# Multiarrangement: A Plug & Play Geometric Data Collection Package For Video Stimuli

**DOI:** 10.1101/2025.10.17.683124

**Authors:** Umur Yıldız, Burcu A. Ürgen

## Abstract

We present Multiarrangement, an offline, open-source Python toolkit for collecting human similarity judgments for video stimuli through multi-arrangement tasks. Participants arrange subsets of stimuli in a 2D arena such that Euclidean distances reflect perceived dissimilarity. The toolkit supports two experimental paradigms: a set-cover scheduling system that uses combinatorial covering designs, aiming to avoid overwhelming the participant for stimulus-rich settings, and an adaptive Lift-the-Weakest scheduler that focuses each new trial on the globally least certain pair and informative neighbors. Across trials, partial distance evidence is fused into a representational dissimilarity matrix. Further refinement is also available with optional reliability-like weighting and inverse MDS which can reduce cross-trial prediction error. We document task design, algorithms, a small within-subject validation, and provide practical guidance for reliable use.

## Introduction

Video has become a first-class stimulus in many behavioral and neural research paradigms. Naturalistic movies capture rich, time-varying perception and cognition and are now common in neuroimaging (Hanke et al., 2014; Liu et al., 2022), where the shared narrative time-locks cortical responses across viewers (Hasson et al., 2004) and yields more reliable, behavior-predictive functional connectivity than rest (Finn et al., 2020). At the behavioral level, validated film-clip corpora reliably elicit target emotions and support ecologically valid laboratory experiments (Gilman et al., 2017; Gross & Levenson, 1995; Schaefer et al., 2010). Naturalistic videos also strengthen behavioral paradigms by preserving multimodal dynamics, narrative structure, and social context (Baldassano et al., 2017; Nastase et al., 2020; Sonkusare et al., 2019). This results in measurements that generalize better to everyday cognition and exposes processes such as event segmentation and predictive processing that are often not well captured by static stimuli (Baldassano et al., 2017; Nastase et al., 2020; Sonkusare et al., 2019). Furthermore, videos differ from their static counterparts in that they carry a manipulable temporal and audiovisual structure. Researchers can reorder shots or remove sound while keeping content realistic, and these manipulations have measurable effects on reliability and cognition (Hasson et al., 2008; Jääskeläinen et al., 2021; Lerner et al., 2011).

Although the value of video stimuli is better appreciated, most video-rating workflows are sequential, where participants watch a clip and provide a rating on a scale. Sequential methods are well matched to scalar targets (MOS, valence), but they are inefficient for capturing relational structure among many clips, and paired-comparison protocols or triplet designs scale poorly (*O*(*N* ^2^) or *O*(*N* ^3^)). Contemporary continuous-annotation toolchains for video provide frame-level valence/arousal traces, yet still yield unidimensional trajectories per pass (Baveye et al., 2015; Koelstra et al., 2012). What is missing for video-centric studies is a method that allows participants to see and compare multiple clips at once, while scaling to dozens of stimuli without overwhelming either the display or the participant. No off-the-shelf, non-commercial software package currently exists to facilitate this type of data collection for video stimuli.

Multiarrangement addresses this gap. Participants arrange small subsets on a two-dimensional canvas, and we aggregate distances across trials into a single representational dissimilarity matrix (Kriegeskorte et al., 2008; Nili et al., 2014). Each placement yields many informative pairwise constraints, which reduces the quadratic burden of exhaustive pairwise ratings while preserving fine-grained structure. Balanced subset schedules give broad coverage with modest trial counts and reduce scale-use and anchoring artifacts that often appear in scalar ratings. For controlled studies, the resulting stimulus space supports targeted sampling, preto post-training comparisons of category structure, and explicit tests of dimensional organization using a single dataset. For cognitive neuroscience the same dissimilarity matrix fits directly into representational similarity analysis and model-based encoding or decoding, enabling comparisons between behavior and brain across images, audio and video, with straightforward reliability checks via split halves and cross-validation. As a library, it provides reproducible scheduling, streamlined interfaces for static and time-varying stimuli, detailed trial logs, and easy integration for downstream analyses.

## Features and Functionality

The design of Multiarrangement focuses on providing a turn-key solution for geometric data collection by combining a straightforward user interface with an efficient back-end for trial management. The goal is to offer a tool that is both accessible for the researcher and effective for data collection. This is accomplished through two main components, which we detail in the following sections:

### User Interface

The user interface of Multiarrangement adheres to a design similar to prior multi-arrangement tools (Kriegeskorte & Mur, 2012), but is tailored for the specific demands of dynamic media. Multiarrangement offers a workspace in which a single circular arrangement area is present. Stimulus tokens initially appear in a seating area outside the circle, from which the user must move them. This two-region design is a deliberate choice, adhering to the paradigm established in seminal multi-arrangement research. The circular workspace is specifically employed to minimize layout biases, as it lacks the corners or implicit axes of a square that might otherwise influence participant placements (Figure 1, panel a). For largesubset trials, an optional, center-locked zoom allows local magnification for precise placement.

**Figure 1.**
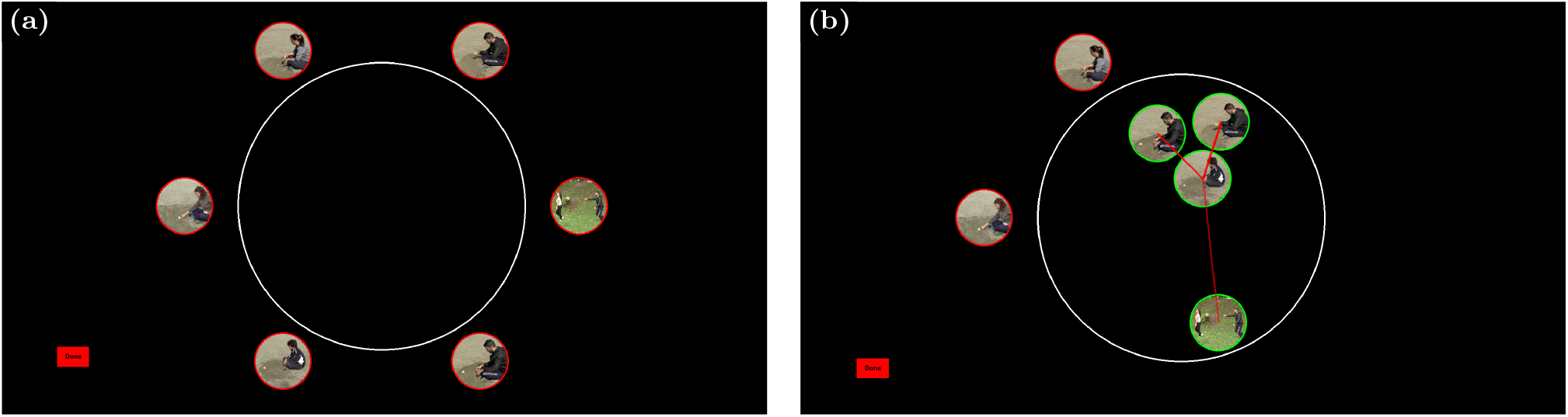
Multiarrangement user interface. The workspace at the start of a trial (a) shows the central circular arrangement area and the peripheral seating area. During placement (b), dragging a token displays transient guidance lines whose thickness/opacity scale with proximity.

**Figure 2.**
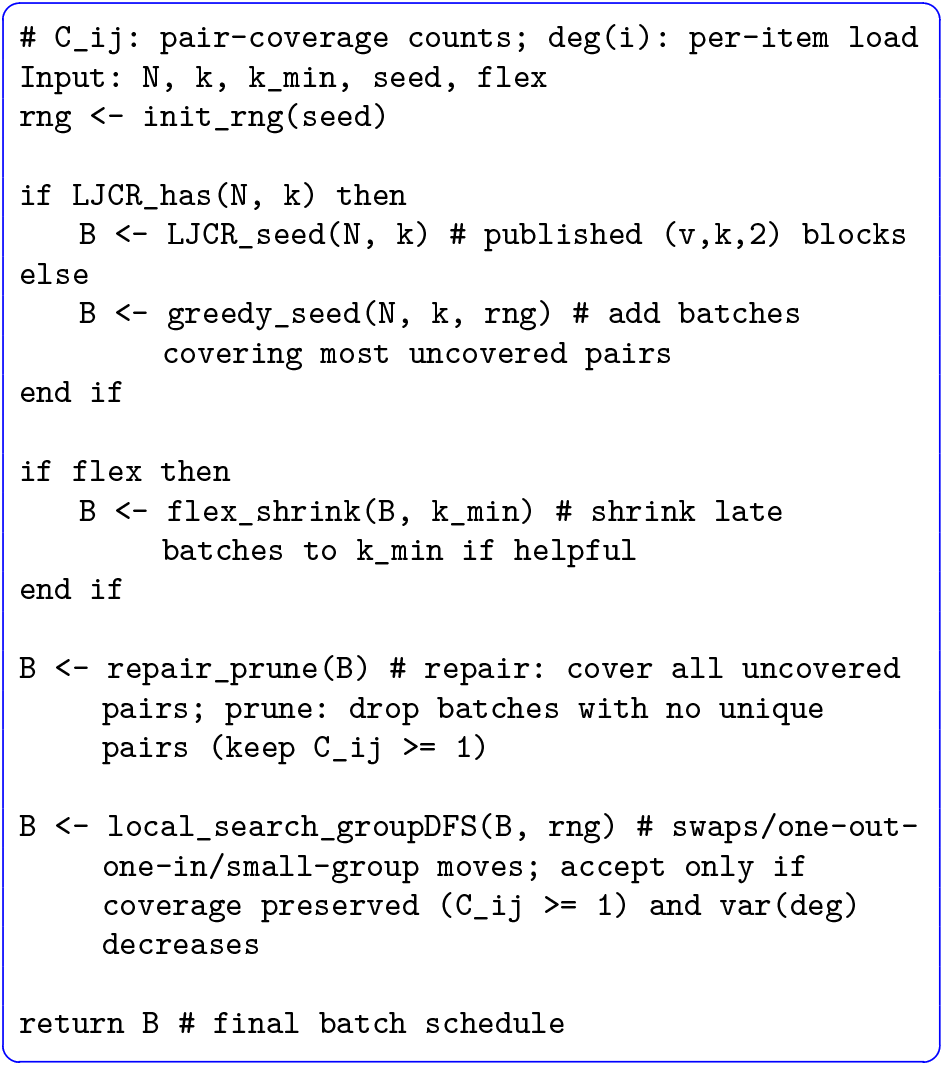
Set-Cover scheduler.^1^ Initialization uses a published (*v, k*, 2) covering when available or a greedy seed otherwise. `flex_shrink` can trim late batches to *k*_min_. `repair_prune` enforces full pair coverage and removes batches contributing no unique pairs, and `local_search_groupDFS` preserves coverage while reducing exposure variance via small swaps/group moves. Returns ordered batches and coverage diagnostics.

The primary mode of interaction is direct manipulation via drag-and-drop. Participants use the mouse to move stimulus tokens (typically thumbnails for videos and images, or icons for audio) from the seating area into the circular workspace. A critical feature, designed specifically for dynamic media, is the ability to inspect stimuli on demand. By double-clicking any stimulus token, the participant can play the associated video in a pop-up window. This allows for repeated review and fine-grained comparisons, a necessary function when judging the similarity of complex, time-varying stimuli.

While a stimulus token is being dragged, transient guidance lines connect it to every other token currently in the circle. Line thickness/opacity scale with proximity, providing a quick visual sense of relative distances (Figure 1, panel b). By default, a trial can be submitted only after every token in the current trial’s subset has been inspected at least once and all tokens are inside the circle.

The interface also uses colored rims around stimulus tokens to signal compliance status: red indicates the requirements for submission pertaining to the specific token have not yet been fulfilled, and green confirms the token satisfies the per-trial requirement. Default instruction sets are provided in Turkish and English, and the interface supports user-specified instructional text and labels via simple templates to enable customization.

## Scheduling Algorithms

The scheduler selects, on each trial, a subset of stimuli that balances simultaneous comparison against per-trial effort while accumulating pairwise evidence toward a reliable RDM (Representational Dissimilarity Matrix) within a fixed session budget. We provide two complementary strategies. First is a SetCover mode that precomputes small batches that together cover all unordered pairs and yields a fixed, between-subject-comparable schedule. In this approach we seed from published (*v, k, t*=2) coverings in the La Jolla Covering Repository (Gordon, 2025; Gordon et al., 1995) and try to provide more balanced batches. The second strategy is an adaptive mode that maintains an evidence matrix, selects the globally weakest pair as anchors, and expands to a bounded subset until an evidence threshold or time limit is reached. We follow the “Lift-the-Weakest” formulation of Kriegeskorte and Mur (2012) with some modifications.

## Set-Cover Implementation

### Problem setup

Let *ℐ* = *{*1, …, *N }* index the stimuli. A trial presents a batch *B* ⊂ *ℐ* of size *k*, covering the intra–batch pairs *𝒫* (*B*) = *{*(*i, j*) : *i < j, i, j* ∈ *B}*. We seek a small family 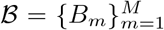 such that 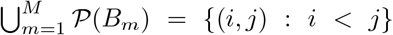 while bounding *k* to avoid on-screen overload and keeping *M* small enough to fit the session budget.

### Construction

When available, we seed from published (*v*=*N, k, t*=2) coverings in the La Jolla Covering Repository, otherwise we initialize greedily by iteratively adding a batch that covers the largest number of yet-uncovered pairs. We then apply a light refinement pipeline consistent with the released code: *repair/prune* to guarantee full coverage while removing batches that add no unique pairs, followed by *local search + group DFS* to smooth per-item load via item swaps and small group moves without breaking coverage. A shrink-only variant (*flex*) may reduce some late batches to *k*_min_ when this closes residual gaps or trims *M* without increasing per-trial burden.

### Execution controls for video

We show one batch per trial. For video, we keep *k* in a narrow range (e.g., 6 ≤ *k* ≤ 10) to bound playback and interaction time, randomize initial token positions on the screen to reduce placement bias, and interleave batches so that no stimulus appears in adjacent trials beyond a user-specified limit. The schedule, RNG seed, and coverage diagnostics are recorded for reproducibility.

### Set-Cover for video

Set-Cover fixes the per-trial set size *k* and, for fixed *k*, reduces total trials relative to naive pairwise enumeration by approximately a factor of 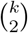. Participants compare a small, consistent number of clips per trial while every unordered pair is co-presented at least once. To control duplication, we refine the schedule to remove redundant batches and balance per-item exposure deg *i* via local swaps and prune & repair iterations, which minimizes repeated co-occurrences and distributes necessary repeats across items and pairs. For remaining repeats, we aggregate distances using either an RMS-matched estimator, which aligns each trial’s scale to the running estimate, a hybrid estimator, which keeps that alignment while giving more weight to clearer separations, or a maximum distance scaled variant, which normalizes each trial by its largest distance for bounded, simple updates. We do not renormalize the global matrix between batches. Each trial is aligned to the current scale using only overlapping already observed pairs. A single final rescaling to unit off-diagonal RMS is applied after all pairs are observed. These estimators also come with optional robust Winsor or Huber reweighting and a subsequent optional inverse MDS step. Together, these steps preserve the time efficiency and manageable on-screen complexity of a fixed-batch design while yielding a smoother, globally consistent RDM.

## Lift-the-Weakest Implementation

### Problem setup

Adaptive scheduling maintains an evidence matrix *W* ∈ ℝ^*N×N*^ that accumulates support for pairwise dissimilarities across trials. At each step the scheduler targets the least supported region while keeping per-trial effort bounded, following the Lift-the-Weakest principle (Kriegeskorte & Mur, 2012).

### Policy

Lift-the-weakest begins with an initial rating step (Trial 0) in which all *N* items are displayed simultaneously. Each subsequent trial anchors on the globally weakest pair (*a, b*) = arg min_*i<j*_ *W*_*ij*_. Starting with *S* = *{a, b }*, items are added greedily up to a size bound to increase expected information, preferring candidates that create low-evidence links to members of *S*. After a layout is collected, the trial is off-diagonal RMS-matched to the current estimate. Weights are computed from the raw on-screen distances and increase quadratically with distance, while the numerator uses the RMS-matched scaled distances. The run stops when the minimum utility *u*(*W*) = −1 exp(−*κW*) exceeds a threshold *u*^*^ or, equivalently, when the minimum evidence threshold is reached (min_*i<j*_ *W*_*ij*_ ≥ *w*^*^ = −ln(1− *u*^*^)/*κ)*, or finally when a trial or time cap is reached (Kriegeskorte & Mur, 2012).

### Execution controls for video

Subset size is bounded *k*_min_ ≤ |*S* | ≤*k*_max_ to keep playback and interaction time practical. Light diversity constraints limit consecutive reuse of the same anchors and discourage high Jaccard overlap between successive subsets. A limited-iteration inverse MDS refinement can be applied intermittently or at the end to improve cross-trial consistency without a large runtime (Kriegeskorte & Mur, 2012).

### Video-specific adaptations

We provide an adaptive Lift-the-Weakest (LtW) mode (Kriegeskorte & Mur, 2012) and introduce small, pragmatic modifications for dynamic video stimuli. Expansion uses a cost-aware utility that divides expected evidence gain by an estimated per-item review cost based on clip duration and recent playback behavior, aiming to keep per-trial time near a target. To avoid starving long clips, we cap the per-item cost estimate and enforce a small minimum inclusion rate for long-duration items. Subset size is chosen within bounds to satisfy a per-trial time target rather than a fixed *k* (i.e., *k*_*t*_ ∈ [*k*_min_, *k*_max_]). Light diversity constraints use soft penalties to limit anchor reuse and discourage excessive Jaccard overlap between successive subsets to reduce fatigue without overriding information gain. Boundary-focused expansion prioritizes candidates that increase evidence around locally uncertain regions by combining low-evidence links with a simple local stress score. Evidence updating supports three modes. The default hybrid mode RMS matches each trial to the current estimate, computes residuals on the RMS scale, and weights pair updates by raw onscreen distance raised to *α*. Winsorization applies to these raw-distance weights and Huber reweighting applies to residuals. The RMS-only mode RMS matches each trial and uses uniform pair weights. Huber reweighting still applies and Winsorization has no effect in this mode. The max-scaled mode rescales each trial so its largest pair spans the layout. It supports either uniform weights or raw-distance weights with the same options. In the hybrid and RMS-only modes we renormalize the RDM after each fuse so that the off-diagonal RMS equals 1. In the max-scaled mode this renormalization is optional and is off by default in order to preserve the interpretation that the largest pair spans the layout. An inverse-MDS update can be applied intermittently or at the end; we use a small, fixed number of iterations in practice to balance runtime and stability. Defaults and exact operationalizations of these parameters (cost model, stress score, diversity penalties) and their usage are documented in our GitHub repository.^2^

## Validation

### Protocol

We compared multi-arrangement RDMs against one-by-one comparison RDMs within-subject (n=2) on the 58-item set of natural human action videos from a validated corpus (Urgen et al., 2023). Multi-arrangement trials followed a fixed Set-Cover schedule with *N* = 58 and batch size *k* = 8, guaranteeing at least one co-presentation of every unordered pair while bounding per-trial load. Fusion used the per-trial scaled estimator with *α* = 0 (max-normalized distances, no distance-based weighting) and inverse-MDS disabled. Convergence was quantified using Pearson and Spearman correlations of the raw off-diagonal entries (vectorized lower triangle; *N* = 58 ⇒ 1,653 pairs). Agreement on the raw scale was assessed with the concordance correlation coefficient (CCC; Lin 1989), Deming regression with variance ratio *λ* = 1 and 95% confidence intervals (Carstensen, 2010; Linnet, 1993), RMSE, and Bland–Altman bias and 95% limits of agreement (Bland & Altman, 1986; Giavarina, 2015). For range-standardized error and visualization only, we additionally report metrics after separate per-method min-max scaling to [0, 1]. Summary metrics are reported in Table 1.

**Table 1.** Agreement between one-by-one and multi-arrangement dissimilarities.

| Correlation and error metrics |  |  |  |  |
| --- | --- | --- | --- | --- |
| Subject | Pearson | Spearman | CCC | RMSE<sub>norm</sub> |
| Subject 1 | 0.846 | 0.744 | 0.778 | 0.207 |
| Subject 2 | 0.694 | 0.593 | 0.578 | 0.200 |
| Mean | 0.770 | 0.669 | 0.678 | 0.203 |

| Deming regression |  |  |  |  |
| --- | --- | --- | --- | --- |
| Subject | Slope | CI<sub>low</sub> | CI<sub>high</sub> | Intercept |
| Subject 1 | 0.934 | 0.914 | 0.956 | −0.073 |
| Subject 2 | 0.872 | 0.827 | 0.911 | −0.024 |

| Bland–Altman |  |  |  |
| --- | --- | --- | --- |
| Subject | Mean diff | LoA<sub>low</sub> | LoA<sub>high</sub> |
| Subject 1 | 0.123 | −0.204 | 0.450 |
| Subject 2 | 0.120 | −0.175 | 0.415 |

### Interpretation

Subject 1 shows high correlation and substantial concordance; Subject 2 shows moderate correlation and concordance. Deming slopes below 1.0 with small negative intercepts indicate that multi-arrangement yields slightly compressed dissimilarities relative to one-by-one overall, consistent with a mild range contraction in simultaneous layouts. Bland–Altman analysis shows a small positive bias for one-by-one over multi-arrangement with reasonably narrow limits of agreement.

Importantly, we do not expect perfect alignment between the two methods. Multi-arrangement elicits context-infused judgments where each item is evaluated relative to the specific subset presented, whereas sequential ratings may invoke implicit comparison against different internal anchors. Nevertheless, the systematic relationship indicates that both methods converge on the similar underlying similarity structure (see Table 1 and Figure 4).

**Figure 3.**
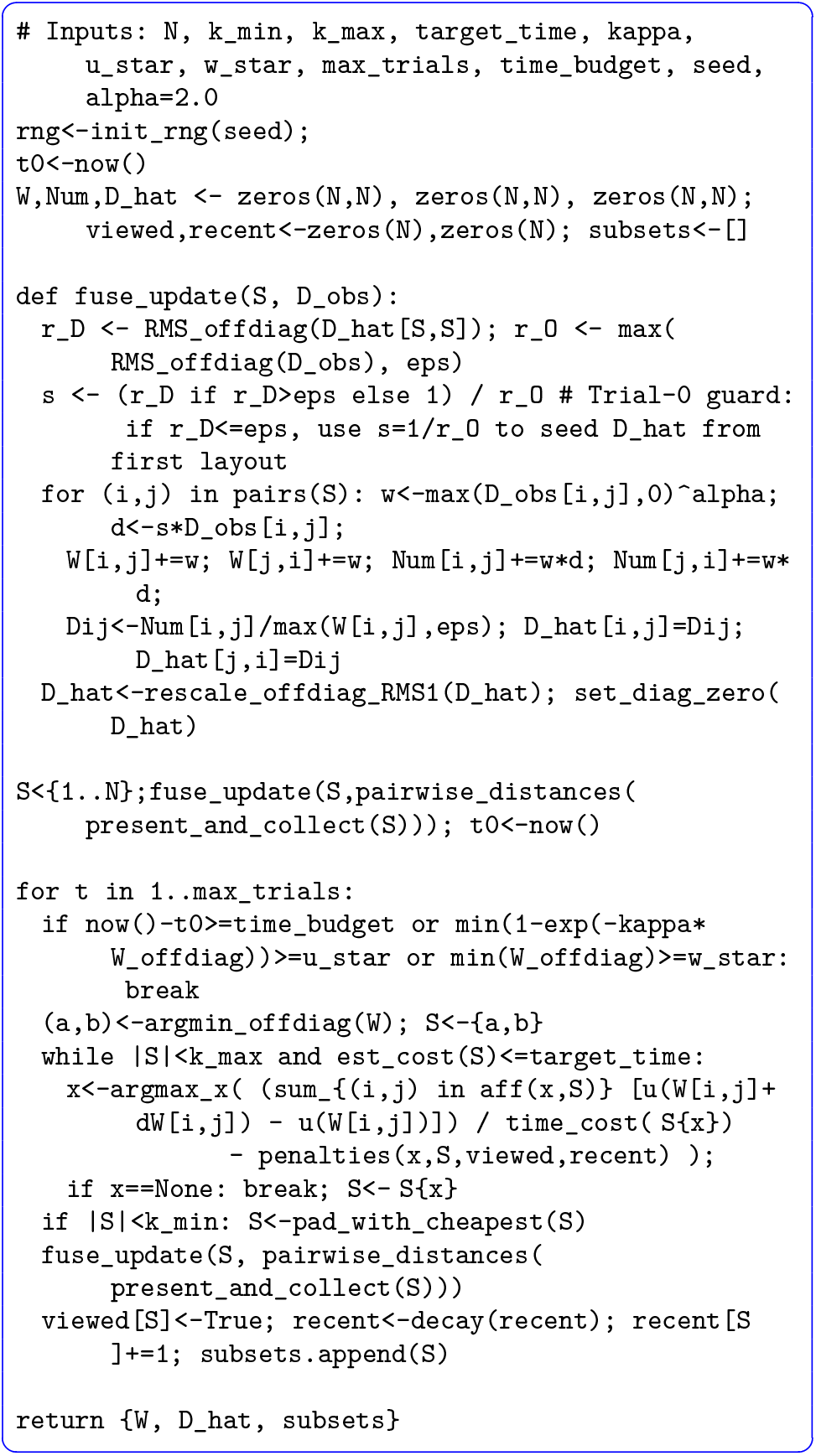
Lift-the-Weakest with initial rating step. Trial 0 shows all *N* items to orient participants. Each subsequent step anchors the current weakest-evidence pair under *W*, grows subset *S* within size/- time bounds to maximize expected gain, collects a layout, RMS-matches to 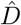, updates *W*, Num, 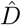 using weights based on raw on-screen distances^*α*^ with the numerator computed from RMS-matched scaled distances, renormalizes to off-diagonal RMS = 1, and stops at utility threshold *u*_*_, minimum evidence threshold *w*_*_, or the time cap.

**Figure 4.**
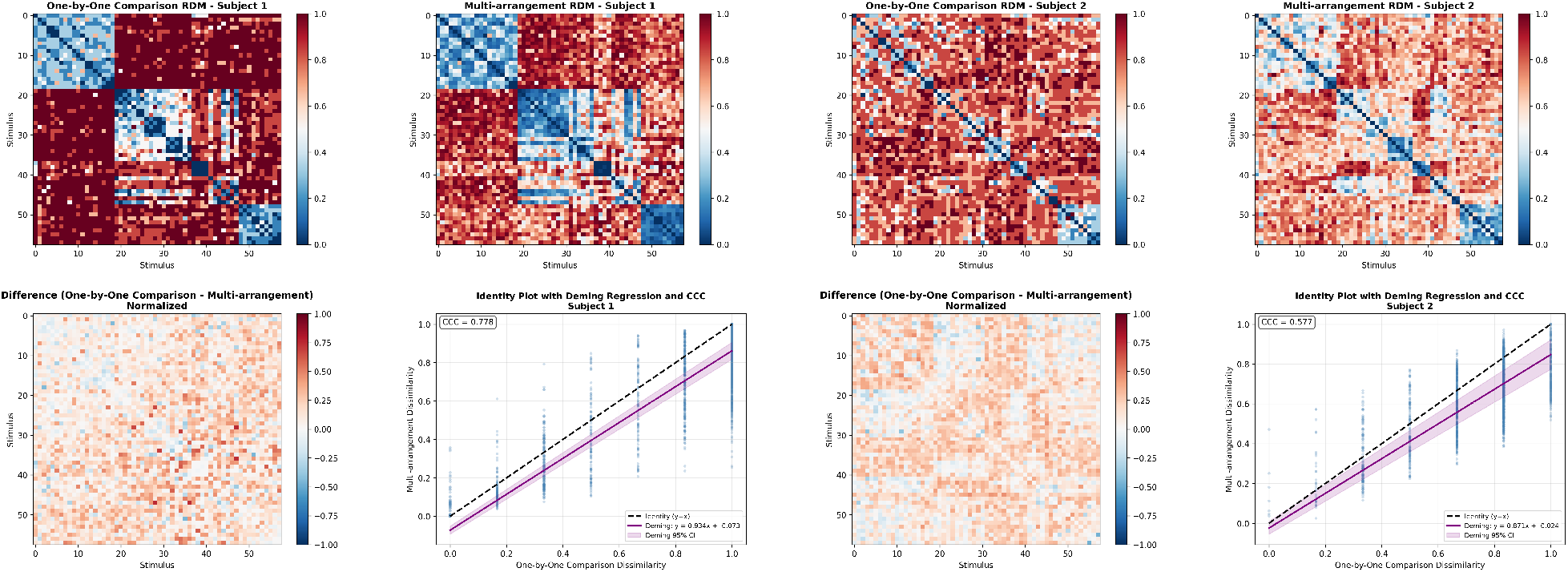
RDM agreement for two subjects. Panels correspond to Subject 1 (left) and Subject 2 (right). Within each panel: top row shows the one-by-one and multi-arrangement RDMs. Bottom left shows the normalized differences (one-by-one minus multi-arrangement) while the bottom right shows the identity plot with Deming regression and CCC.

## Usage of the Package

### Installation and setup

Install from PyPI (Python 3.12+) and ensure common multimedia dependencies are available.

~~~
pip install multiarrangement
# or: uv pip install multiarrangement
~~~

The package bundles demo media (videos, images, audio), default instruction clips, and a cache of covering designs (LJCR) for offline use.

### Preparing stimuli

Place your files (videos, audio, or images) in a single input folder. The type of modality is detected automatically, and requirements for submissions are adaptively set, and mixed modalities are supported. Filenames serve as token labels for easy tracking of entries.

### Fixed-batch (Set-Cover) Usage

The following example demonstrates how to use the Set-Cover mode in our API. Firstly we precompute a fixed-*k* schedule of covering batches, seeding for reproducibility and using La Jolla coverings when available, then launch the session with that schedule to collect placements and automatically save the resulting RDM and logs. Batch size *k* can be tuned and shrink-only refinement can be enabled with `flex=True`. Weight modes and alphas can be tuned as parameters based on preferences and assumptions. The run writes per-trial logs, coverage diagnostics, and the final RDM to the output folder for use in visualization or further downstream analysis.

~~~
import multiarrangement as ma
input_dir = “./videos” # your stimuli
output_dir = “./results”
batches = ma.create_batches(
  ma.auto_detect_stimuli(input_dir),
  k=8, seed=42, flex=False
)
results = ma.multiarrangement(
  input_dir=“./videos”, #Where your videos or audios
     are
  batches=batches,
  output_dir=“./results”, #Where your results will
     *appear*
  show_first_frames=True,
  fullscreen=False,
  language=“en”, # Or tr if you’d like Turkish
    *instructions*
  instructions=“default”, # or None, or [“Custom”, “
     lines”]
 setcover_weight_alpha=2.0,
 setcover_weight_mode=‘max’, # ‘max’ (d/max) or ‘
     rms’ (‐RMSmatched) or ‘k2012’ for hybrid LtW
    *style*
 use_inverse_mds=False,
 robust_method=None, # ‘winsor’ or ‘huber’
)
 results.vis(title=“‐SetCover RDM”)
 results.savefig(“results/rdm_setcover.png”, title=“
  ‐SetCover RDM”)
~~~

### Adaptive (Lift-the-Weakest) Usage

The following example demonstrates how to run the adaptive Lift-the-Weakest mode within the Multiarrangement library: set subset size bounds and a target per-trial time or evidence threshold, optionally enable inverse MDS, then run to produce the evolving RDM, evidence matrix, and logs. The LtW approach is more dynamic and fine-grained than Set-Cover in how it revisits stimulus pairings, but can take longer and may overwhelm participants when N is large, both in total time and during the initial arrangement phase.

~~~
import multiarrangement as ma
res = ma.multiarrangement_adaptive(
  input_dir=“./videos”,
  output_dir=“./results”,
  participant_id=“S01”,
  language=“en”,
  fullscreen=True,
  min_subset_size=4, max_subset_size=6,
  evidence_weight_mode=“k2012”, # “rms” or “max”
  evidence_alpha=2.0,
  stop_on_utility=False, # use raw-evidence
   *threshold*
  evidence_threshold=0.35,
  instructions=“default”,
 )
  res.vis(title=“Adaptive LTW RDM”)
  res.savefig(“./results/rdm_adaptive.png”, title=“
   Adaptive LTW RDM”)
~~~

### Custom instructions and localization

Default instructions are provided in English and Turkish for convenience. To override, pass a list of strings to be shown as paginated screens before the task, per the following example:

~~~
custom = [
  “Welcome to the study.”,
  “Drag each item into the white circle.”,
  “Double-click a token to play/replay the video.”,
  “Press SPACE to continue.”
]
ma.multiarrangement(input_dir, batches, “./results”,
   instructions=custom)
# ...or for adaptive:
ma.multiarrangement_adaptive(“./videos”, “./results”,
   instructions=custom)
~~~

For further detailed documentation, see our GitHub page^2^. We release all code, experiment templates, and analysis scripts used in this paper.

## Conclusions

In this paper, we introduce Multiarrangement, a turn-key, offline, open-source software package for collecting human similarity judgments with dynamic visual stimuli. The system unifies a simple, robust interface with a circular arena, on-demand playback, compliance cues, optional zoom, and bilingual defaults with scheduling back-ends that make large stimulus sets practical. A fixed Set-Cover mode produces deterministic, between-subject-comparable schedules with bounded per-trial load, while an adaptive Lift-the-Weakest mode targets the least-supported region of the space under explicit time and subset-size constraints. Both schedulers offer choices for distance fusion, stopping rules, and robustness & refinement, allowing researchers to align the pipeline with their design goals and assumptions. We also report a small within-subject validation indicating that multiarrangement approximates the structure captured by one-by-one comparisons while using far fewer trials, enabling context-aware judgments at lower participant burden.

Beyond video, the same workflow supports images and audio with no changes to the analysis path, and results are written in standard formats to facilitate downstream use in behavioral and neuroimaging pipelines. The package is designed to be easily extended. Researchers can customize instructional text, enforce additional compliance rules, adjust scheduler parameters, and integrate new qualitycontrol or weighting schemes while preserving reproducibility via recorded seeds and schedules. We hope this plug-and-play framework lowers the barrier to deploying multi-arrangement studies at scale and stimulates new experimental designs that leverage simultaneous comparison to probe perceptual and cognitive structure in richer, more naturalistic settings.

## Declarations

### Authors’ contributions

Both authors jointly conceptualized the study. U.Y. implemented the software, conducted testing & validation, and collected the data. Both authors co-wrote the first draft, critically revised subsequent drafts, and approved the final manuscript for submission.

### Funding

This research received no specific grant from any funding agency, commercial or not-for-profit sectors.

### Competing interests

The authors declare no competing interests.

### Ethics approval

The study was approved by, and conducted along the guidelines of, the Bilkent University Ethics Committee.

### Consent to participate

All participants provided written informed consent prior to participation. Data were analyzed anonymously.

### Consent for publication

All participants provided written informed consent for publication of anonymized data.

### Open Practices Statement

Materials (stimulus lists and templates), validation data, and analysis code are available at 10.5281/zenodo.17463843. Further details on package use are available in our GitHub repository at github.com/UYildiz12/Multiarrangementfor-videos. None of the reported studies were preregistered.

## Footnotes

1 Algorithm pseudocode listings are framed in blue, and runnable code examples are framed in black.

2 https://github.com/UYildiz12/Multiarrangement-for-videos

## Notes

https://github.com/UYildiz12/Multiarrangement-for-videos

### Competing Interest Statement

The authors have declared no competing interest.

### Summary of Updates

Details on pseudocode, and updates on library usage.

